# Predicted effects of severing enzymes on the length distribution and total mass of microtubules

**DOI:** 10.1101/752006

**Authors:** Y.-W. Kuo, O. Trottier, J. Howard

## Abstract

Microtubules are dynamic cytoskeletal polymers whose growth and shrinkage are highly regulated as eukaryotic cells change shape, move and divide. One family of microtubule regulators includes the ATP-hydrolyzing enzymes spastin, katanin and fidgetin, which sever microtubule polymers into shorter fragments. Paradoxically, severases can increase microtubule number and mass in cells. Recent work with purified spastin and katanin accounts for this phenotype by showing that, in addition to severing, these enzymes modulate microtubule dynamics by accelerating the conversion of microtubules to the growing state and thereby promoting their regrowth. This leads to the observed exponential increase in microtubule mass. Spastin also influences the steady-state distribution of microtubule lengths, changing it from an exponential, as predicted by models of microtubule dynamic instability, to a peaked distribution. This effect of severing and regrowth by spastin on the microtubule length distribution has not been explained theoretically. To solve this problem, we formulated and solved a master equation for the time evolution of microtubule lengths in the presence of severing and microtubule dynamic instability. We then obtained numerical solutions to the steady-state length distribution and showed that the rate of severing and the speed of microtubule growth are the dominant parameters determining the steady-state length distribution. Furthermore, we found that the amplification rate is predicted to increase with severing, which is a new result. Our results establish a theoretical basis for how severing and dynamics together can serve to nucleate new microtubules, constituting a versatile mechanism to regulate microtubule length and mass.

**Significance:** The numbers and lengths of microtubules are tightly regulated in cells. Severing enzymes fragment microtubules into shorter filaments and are important for cell division and tissue development. Previous work has shown that severing can lead to an increase in total microtubule number and mass, but the effect of severing on microtubule length is not understood quantitatively. Combining mathematical modeling and computational simulation, we solve the microtubule length distribution in the presence of severing enzymes and explore how severing activity and microtubule dynamics collectively control microtubule number and length. These results advance our understanding of the physical basis of severing as a regulatory mechanism shaping the cellular cytoskeletal network.

## Introduction

The cytoskeleton is a network of filamentous polymers and is found in all living organisms including bacteria, plants and animals. Microtubules form one component of the eukaryotic cytoskeleton. They elongate from their ends by addition of tubulin subunits, and alternate between phases of slow growth and rapid shrinkage. This alternation, termed dynamic instability, causes frequent turnover of polymer and exchange of tubulin subunits with the soluble pool (1, 2). The dynamics of microtubules can be described quantitatively by four parameters: the growth (polymerization) rate, the shrinkage (depolymerization) rate, the catastrophe frequency (the transition from growing to shrinking states) and the rescue frequency (the transition from shrinking to growing states)(2, 3). As eukaryotic cells undergo cell division, migration or shape change, microtubule-associated proteins (MAPs) regulate the dynamics of microtubules so as to alter their numbers and lengths (4). Much is known about the mechanisms by which MAPs nucleate microtubules, accelerate growth, promote or inhibit catastrophe, increase rescue or induce depolymerization (5, 6). However, the mechanisms by which microtubule severing enzymes regulate the microtubule cytoskeleton is not well understood.

The microtubule severing enzymes spastin, katanin and fidgetin are AAA-ATPases that use the chemical energy of ATP-hydrolysis to sever microtubules into shorter filaments by generating internal breaks in the microtubule lattice(7–9). Microtubule severing, first observed in *X. laevis* oocyte extracts(10), was initially thought of as a destructive process when the first severing enzyme katanin was discovered(7). And indeed, when katanin and spastin are overexpressed in tissue culture cells, the amount of microtubule mass is reduced(11–13). However, *in vivo* experiments showed, paradoxically, that genetic knockdown and mutations of severases actually reduce microtubule mass in neurons of flies and fish(14–16) and in the meiotic spindle of worms(17), and reduce the growth of longitudinal cortical microtubule arrays in plants(18). These observations suggest that severases have a nucleation-like activity.

Recent *in vitro* studies have demonstrated that spastin and katanin indeed possess a nucleation-like activities(19, 20). They promote the regrowth of severed microtubules by increasing the frequency of rescue and decreasing the shrinkage velocity and thus lead to a net increase in microtubule number and mass. An extension of the Dogterom & Leibler dynamic instability model(21) to include severing successfully predicted the exponential amplification of total microtubule mass observed *in vitro* and confirmed that the modulation of dynamics is essential to increase microtubule number and mass(19). Spastin-mediated severing also changes the length distribution of microtubules from a monotonically decreasing function to a peaked function(19). The theoretical basis of this effect, which makes the microtubules more uniformly distributed in length, is not understood.

Here we solve an extended dynamic instability model to investigate the microtubule length distribution when the microtubule number and mass are amplified by the combination of severing and regrowth. Our work builds on earlier models for actin (22–25) where severing is necessary to keep the mean filament length finite when the actin concentration is well above the critical concentration for growth. Severing acts as a negative feedback on length because longer filaments are cut more frequently. A similar argument holds for microtubules. However, dynamic instability(21) leads to more complex microtubule behaviors compared to actin, which does not undergo dynamic instability. A theoretical model by Tindemans & Mulder solved the case where the microtubule number is constant(26). In this paper, we solve the Tindemans & Mulder model in the case where the microtubule number and total polymer mass increase, as is observed *in vitro* and in cells.

## Materials and Methods

### Time evolution of microtubule lengths in the presence of severing and dynamics

In a recent paper(19), we solved a generalization of the dynamic instability model(21) that includes microtubule severing(26):

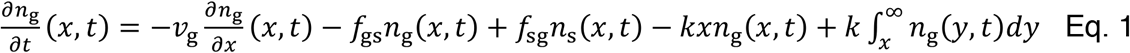

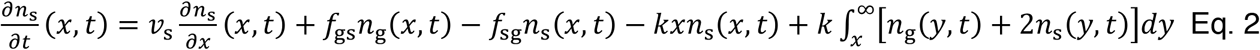

The number of growing and shrinking microtubule plus ends of length *x* at time *t* are denoted by *n*_*g*_(*x*, *t*) and *n*_*s*_(*x*, *t*) respectively. The four dynamic parameters are represented by *v*_*g*_, *v*_*s*_, *f*_*gs*_, *f*_*sg*_: the growth rate, shrinkage rate, catastrophe frequency and rescue frequency, respectively.

In Eq. 1 and 2, the first three terms originate from dynamic instability. The penultimate term represents the disappearance of microtubules of length *x* due to severing (with severing rate *k*). The last term represents severing of microtubules of length greater than *x* into two fragments, one of which has length *x*.

The model makes several additional assumptions: (i) The new plus end is shrinking, while the new minus end is stable. This is based on the observation that around 80% of new ends satisfy this property ((18, 19, 27, 28), but see Vemu et al. 2018 who reported a lower percentage, though this might be due to rapid rescues giving rise to apparent growing ends (20)). A non-zero probability that newly created plus ends grow can be included within the same framework and will be discussed later. (ii) Microtubule dynamics is dominated by the plus ends. In other words, minus ends are considered “passive” in the sense that they neither grow nor shrink, though a minus end can disappear when the plus end depolymerizes all the way back to the minus end. This assumption is based on the reduced dynamics of minus ends, which are often capped or anchored *in vivo* (5, 29). The model could be extended by also considering minus end dynamics, but we have omitted this for the sake of simplicity. (iii) Severing is an instantaneous event that takes place stochastically at a random location on the microtubule lattice with uniform probability. This has experimental support: the location of spastin severing events on microtubules is consistent with a uniform distribution (Fig. S1). The severing rate *k* is constant and has units of length^−1^⋅time^−1^. These assumptions are identical to those made by Tindemans and Mulder.

We solve the equations with following boundary condition:

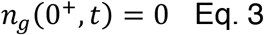

which corresponds to the absence of stable seeds and without spontaneous nucleation. This boundary condition differs from the case solved by Tindemans and Mulder who assumed a constant nucleation rate. In addition, unlike Tindemans and Mulder, we solve for the case where the number of microtubules is increasing (which corresponds to the unbounded growth regime of the Dogterom & Leibler model).

### Computational simulation of the stochastic severing model

To verify the existence of a length distribution at steady-state, we simulated the stochastic equation for microtubule dynamics that includes severing (Eq. 1 and Eq. 2) together with the boundary condition (Eq. 3). The simulation starts with 100 microtubules whose lengths are sampled randomly from an exponential distribution, which is motivated by the steady-state solution of the Dogterom & Leibler dynamic instability model(21). At each time step, the length and state (growing or shrinking) of microtubules may change. In the growing state, a microtubule either grows, undergoes catastrophe and becomes a shrinking microtubule, or is severed and becomes shorter. Similarly, in the shrinking state, a microtubule either shrinks, undergoes rescue and becomes a growing microtubule, or is severed and becomes shorter. In addition, a microtubule disappears when its plus end shrinks to its minus end (*x* = 0) and each severing event creates an additional shrinking microtubule. Owing to severing and the absence of stable seeds, the total number of microtubules is not constant and the length probability distribution is renormalized at every time point. The model’s input parameters, which we refer to as the dynamic parameters, are growth rate *v*_*g*_, shrinkage rate *v*_*s*_, catastrophe frequency *f*_*gs*_, rescue frequency *f*_*sg*_ and severing rate *k*. The dynamic parameters in the unbounded growth regime were obtained from *in vitro* experiments summarized in Table 1(19). The severing activity *k* is set to 0.05 μm^−1^⋅min^−1^, though the value does not qualitatively affect the existence of a steady-state.

**Table 1.** Summary of the dynamic parameters used in the model. The values used in the mathematical model are from previous experimental measurements (19). For testing the effect of microtubule dynamics, a severing rate of 0.05 μm^−1^⋅min^−1^ was used.

| Model parameters | Input values |
| --- | --- |
| Growth rate $v_g$ | 0.79 ($\mu\text{m}/\text{min}$) |
| Shrinkage rate $v_s$ | 9.9 ($\mu\text{m}/\text{min}$) |
| Catastrophe frequency $f_{gs}$ | 0.098 ($\text{min}^{-1}$) |
| Rescue frequency $f_{sg}$ | 3.12 ($\text{min}^{-1}$) |
| Severing rate $k$ | 0.001-0.25 ($\mu\text{m}^{-1} \cdot \text{min}^{-1}$) |

### Steady-state length distribution and rate of microtubule number and mass increase

When the length distribution reaches a steady-state, we showed previously that the above model predicts that the total number *N* and mass *M* of microtubules increase exponentially with time:

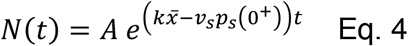

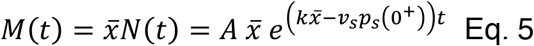

where *A* is a positive constant, *p*_*s*_ is the probability density function of shrinking microtubules and 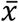 refers to the mean length at steady-state(19). The rate constant inside the exponential in Eq. 4 corresponds to the net creation of new microtubules: it is the difference between the increase in microtubules due to severing 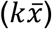 and the decrease due to minus ends disappearing (*v*_*s*_*p*_*s*_(0^+^)).

These equations are a consequence of the assumption that the length distribution of growing microtubules, *p*_*g*_(*x*, *t*), reaches a steady-state and so satisfies:

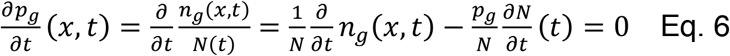

Inserting Eq. 1 and Eq. 4 into Eq. 6 gives the steady-state equation for the growing microtubule distribution:

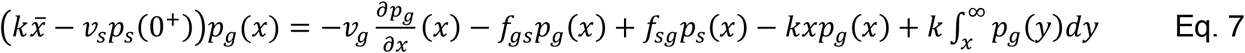

Similarly, the steady-state equation for the shrinking microtubule distribution *p*_*s*_ is:

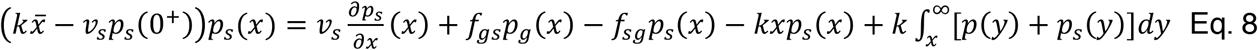

Summing Eq. 7 and Eq. 8, multiplying both sides by *x* and integrating from 0 to infinity:

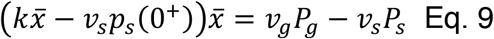

where *P*_*g*_ and *P*_*s*_ are the percentage of growing and shrinking microtubules, respectively. Using Eq. 9 and the fact that *P*_*g*_ and *P*_*s*_ sum to 1, we get:

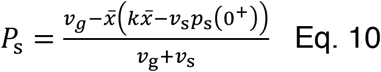

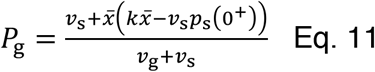

Integrating Eq. 7 with respect to *x* from 0 to infinity:

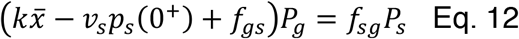

Combining Eq. 10 to 12, we obtain a characteristic equation (the same form as previously derived in(19)):

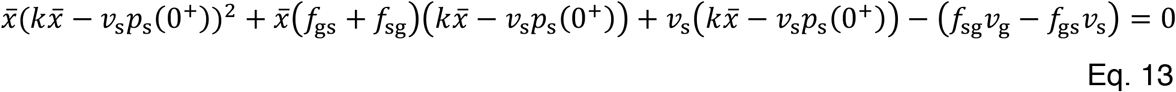

which is a cubic function of the mean length 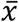 and a quadratic function of *p*_*s*_(0^+^).

The necessary and sufficient condition to find a positive root for 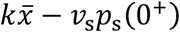, corresponding to the case where the total mass and number of microtubule increase, is that the term *f*_*sg*_*v*_*g*_ − *f*_*gs*_*v*_*s*_ is positive. This is the unbounded growth regime found in the Dogterom & Leibler model where the mean length of microtubules increases indefinitely in the presence of stable microtubule seeds(21). Another way of stating this condition is that *v*_*g*_/*f*_*gs*_ > *v*_*s*_*f*_*sg*_: the mean length increase in the growing state is longer than the mean length decrease in the shrinking state.

### Numerical integration of the steady-state differential equations

Taking derivatives with respect to *x* of the steady-state equations Eq. 7 and Eq. 8 yields a system of coupled second-order ordinary differential equations (ODEs):

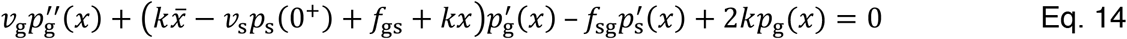

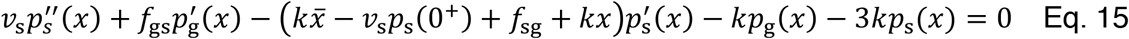

Evaluating Eq. 7 and Eq. 8 at *x* = 0 gives the boundary conditions of the first derivatives:

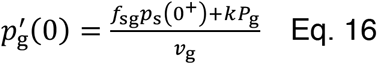

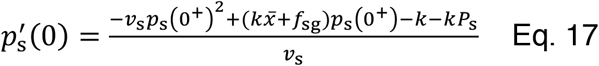

Recall that Eq. 13 is a quadratic equation of *p*_*s*_(0^+^) with the root:

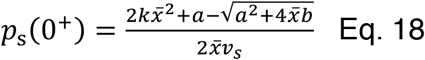

where

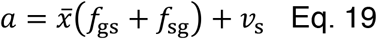

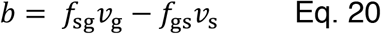

This is the only root of *p*_*s*_(0^+^) that gives a positive value for 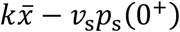, which corresponds to the condition that the total mass and number increases over time with an amplification rate of:

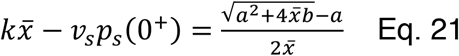

Combining these results, the boundary conditions become:

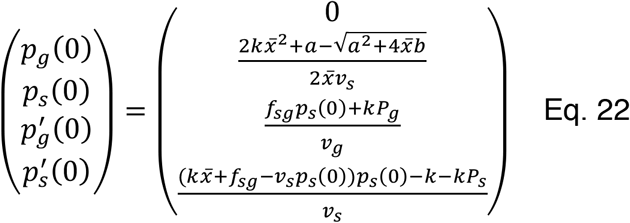

Using the characteristic equation, this set of boundary conditions can be expressed as a function of 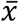, the mean microtubule length, which is the only unknown parameter. The steady-state length distribution in the severing model with dynamic instability is computed numerically with MATLAB using *ode15s*. To integrate the solution, the boundary conditions are evaluated using an estimate for 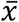 that is iteratively fine-tuned. When the 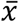 estimate deviates from the true mean length, the solution diverges and the divergence direction depends on whether the input 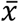 is larger or smaller than the true mean length (see example in Fig. S2A; the blue and red curves diverge in opposite directions). Subsequently, a new estimate for 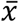 is calculated based on the divergence observed in the previous iteration. Specifically, the 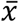 estimate is reduced when a positive divergence was observed (and vice versa). This iteration process terminates when the 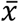 estimate and the mean length calculated from the solution deviate by less than 0.001% (scheme in Fig. S2B). The final solution converges with a pointwise accuracy of at least 10^−5^ μm^−1^. The parameters of the numerical solution are based on the experimentally measured values (see Table 1). To explore the effect of dynamics and severing, each single parameter was altered sequentially.

As a generalization of the model, we allowed newly generated plus ends to be in the growing state, with probability *q*. In the model above, *q* = 0. The generalizations of the master equations (Eq. 1 and Eq. 2) are:

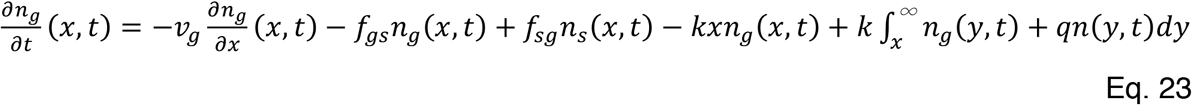

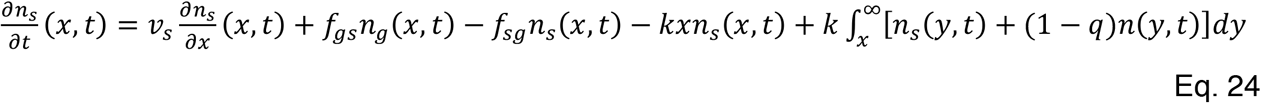

The sum of Eq. 23 and Eq. 24 is identical to the sum of Eq. 1 and Eq. 2 and is independent of *q*. At the length distribution steady-state, the microtubule number and mass increase exponentially with time (Eq. 4 and Eq. 5). Following similar methods described above, we can derive the steady-state length distribution equations as:

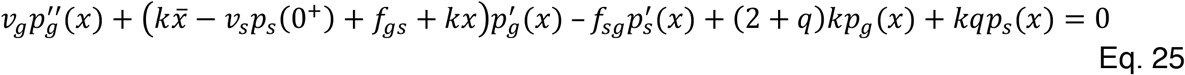

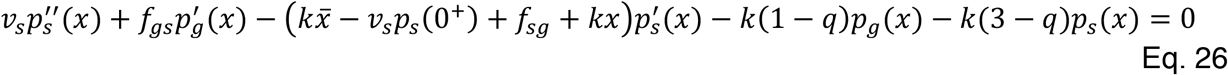

The boundary conditions for these coupled ODEs can also be derived as a function of the mean length 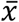:

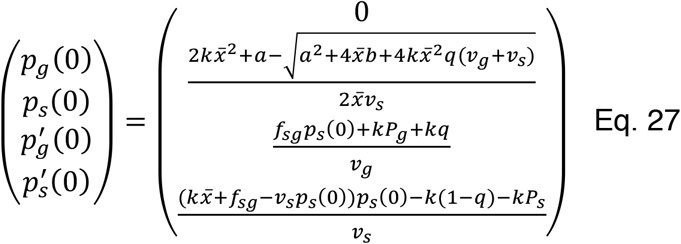

The steady-state length distribution then can be obtained by solving Eq. 25 and Eq. 26 numerically with the boundary condition (Eq. 27), using the aforementioned iteration method.

### Microtubule severing assay

Bovine brain tubulin was purified as previously described(30). Stabilized microtubules were prepared by polymerizing unlabeled tubulin with the slowly hydrolysable GTP analog GMP-CPP (Jena bioscience) and affixed onto the flow channel surface with anti-tubulin antibody (clone SAP.4G5, Sigma Aldrich) following the previous method (31). *Drosophila* spastin was expressed and purified as previously described (19). Severing of the GMP-CPP-stabilized microtubules was performed with 3.5 nM spastin and visualized by interference reflection microscopy (IRM) (32) with a frame rate of 0.5 Hz. Imaging buffer consists of 80 mM K-PIPES, pH 6.9, 1 mM MgCl_2_, 1 mM EGTA, 50 mM KCl supplemented with 1 mM MgATP and 5 mM dithiothreitol. Analysis of severing position was done using Fiji software(33).

## Results and Discussion

### Existence of the length distribution steady-state

To verify that a steady-state length distribution exists, we performed a stochastic simulation of the microtubule severing system (Eq. 1 and Eq. 2, with boundary condition Eq. 3). The initial lengths of microtubules were randomly sampled from an exponential distribution with an average length of 5 μm (Fig. 1A and 1B, blue curves). The initial proportion of shrinking microtubules was set to 10%, though we found that the system still reached the steady-state regardless of this proportion. The dynamic parameters that were used in the simulation are based on previous experimental measurements (Table 1, severing rate *k* = 0.05 μm^−1^⋅min^−1^). The parameters lie in the unbounded growth regime, meaning that in the absence of severing, the mean microtubule length would increase indefinitely.

**Figure 1.**
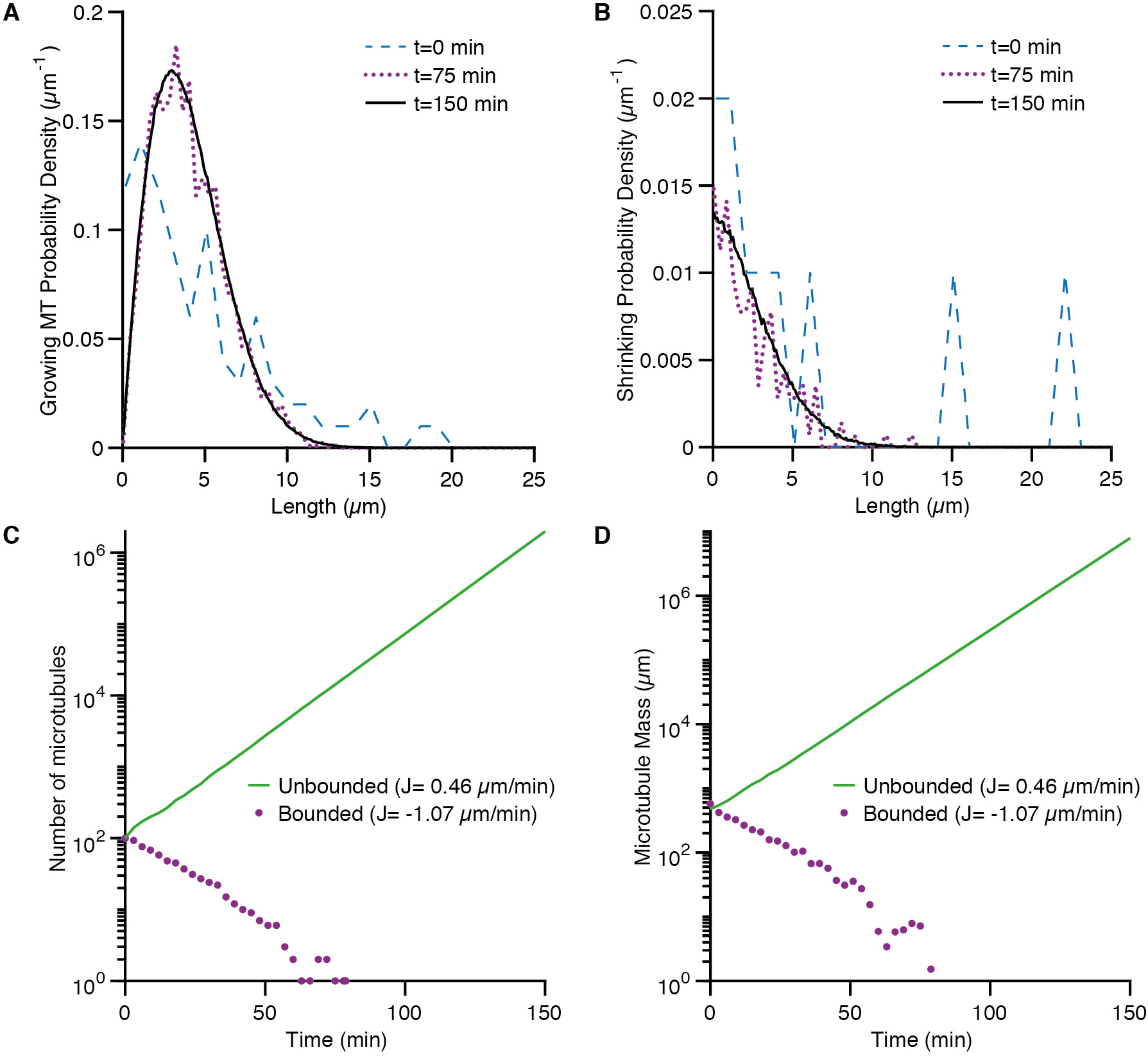
Stochastic simulation of microtubule severing. **A**,**B** Examples of stochastic simulations showing the temporal evolution of the length distributions of growing (left) and shrinking (right) microtubules. **C**,**D** The total number of microtubules (left) and total microtubule mass (right) evolving over time. In the case of unbounded growth, (parameters from Table 1 with *k* = 0.05 μm^−1^⋅min^−1^), both the total number and the mass of microtubules increase exponentially (green solid curve, in the semi-log plots). In the case of bounded growth (parameters from Table 1 but the shrinkage rate is increased to 20 μm/min and the rescue frequency is decreased to 1 min^−1^), the microtubules eventually disappear (magenta dotted curves). The average flux of tubulin onto each microtubule, *J*, is equal to (*f*_*sg*_*v*_*g*_ − *f*_*gs*_*v*_*s*_)/(*f*_*sg*_ + *f*_*gs*_).

The microtubule length distribution converged to a steady-state, which is peaked and has a decaying tail (Fig. 1A). The steady-state shrinking microtubule distribution has a small fraction of zero-length microtubules, *p*_*s*_ 0 (Fig. 1B); the disappearance rate of these shrinking microtubules and the creation rate of new microtubules by severing reach a constant ratio. At steady-state, the total number and mass of microtubules increased exponentially (green solid lines in Fig. 1C and 1D), as predicted by the ordinary differential equations (Eq. 14. and Eq. 15) and the characteristic equation (Eq. 13). Furthermore, when we increased the shrinkage rate and decreased the rescue frequency to enter the bounded growth regime, where *f*_*sg*_*v*_*g*_ − *f*_*gs*_*v*_*s*_ is negative, the number of microtubules went to zero (Fig.1C and Fig. 1D, magenta dotted curves). Thus, the stochastic model confirms the existence of a steady state and that the unbounded growth condition is an essential criterion for amplifying microtubule arrays with severing.

### Simplified no-catastrophe model

In the stochastic simulation, the total number of growing microtubules is much greater than the number of shrinking ones. This implies that, on average, microtubules spend most of their time in the growth phase. This inspired us to consider a simplified case where the microtubules exist only in the growing state, with no catastrophe or shrinkage events. The approximate solution is valid when the effect of dynamic instability is relatively small compared to the net growth, where the tip dynamics can be approximated as a pure drift process with a small diffusion coefficient.

The time evolution of this simplified model can be expressed as:

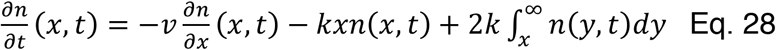

where *v* is the microtubule elongation rate. This equation follows from Eq. 1 with *f*_*sg*_ = 0, *f*_*gs*_ = 0, *n* = *n*_*g*_, *n*_*s*_ = 0, *v*_*g*_ = *v* and we assume that a new plus end is in the growth phase. This equation is similar to the integro-differential equation in Edelstein-Keshet & Ermentrout(22) with the difference that the factor of 2 in the last term of Eq. 23 is replaced by 1, since only one fragment generated from severing is taken into account in their model. With the finite-length assumption, integrating Eq. 28 with respect to *x* from 0 to infinity and combining with the boundary condition *n*(0, *t*) = 0, we get:

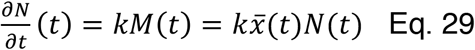

When the length distribution is in the steady-state, the total number and mass of microtubules increase exponentially with a characteristic time 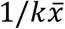. The equation for the length distribution, *p*(*x*), is:

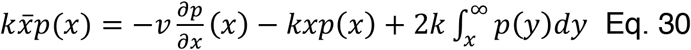

Multiplying by *x* and integrating from 0 to infinity, we find the following expression for the mean length of the distribution:

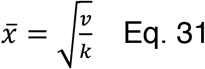

Eq. 30 can be solved analytically (see Appendix) by rewriting it as a Hermite differential equation, which is seen in various physical systems such as the quantum harmonic oscillator, used to model the vibrations of chemical bonds(34). The final solution is:

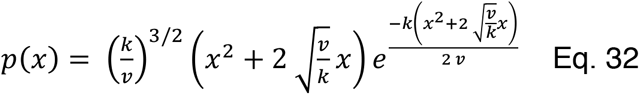

The solution is plotted in Fig. 2: the curve is peaked, starts at 0 when *x* = 0 and decays approximately like a Gaussian at large *x*. Note that the solution is dependent solely on a single parameter: the ratio of elongation rate to severing activity *v*/*k*. With increasing *v*/*k*, the distribution shifts rightwards (Fig. 2).

**Figure 2.**
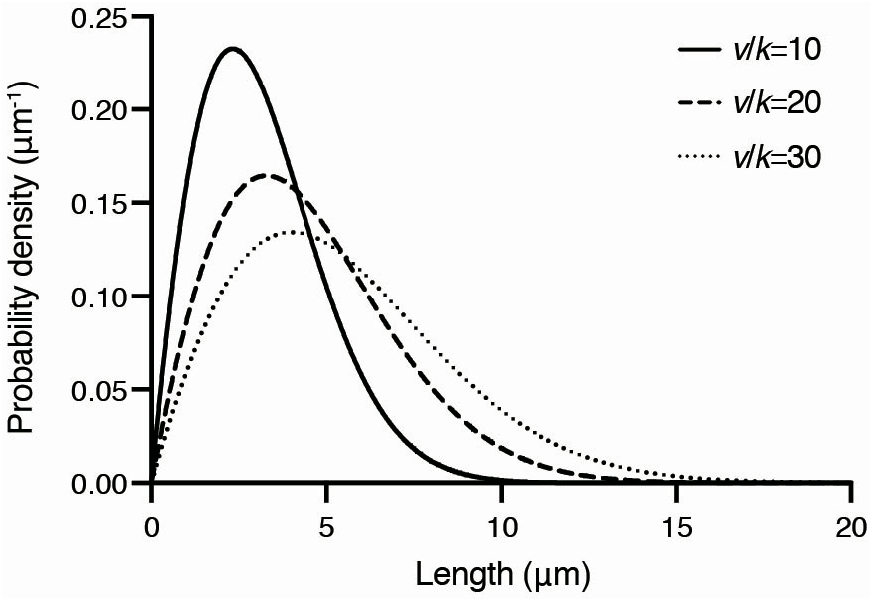
Length distribution of the no-catastrophe model. The analytic solution to the steady-state length distribution (Eq. 32) is plotted for three different values of the ratio of the growth rate to severing rate, (*v*/*k*). As the ratio increases, the lengths increase and the distribution widens.

An important result is that the no-catastrophe model always predicts a finite mean length (provided that *k* > 0). This is explained by the fact that the severing probability increases with length. Therefore, long microtubules are quickly shortened by severing while short microtubules can grow longer before they are severed. This principle also applies to the scenario with dynamic instability because the rate to sever a single microtubule will still increase with polymer length, even in the presence of shrinkage events.

### Numerical solution of the dynamic instability with severing model

To understand the effect of dynamic instability in the presence of severing, we solved the steady-state length distribution numerically (Eq. 14 and 15, see Methods). The growing, shrinking and total microtubule length distributions are shown in Fig 3A-3C. The total microtubule length distribution (Fig. 3C) is mainly determined by the growing microtubules (Fig. 3A), consistent with the stochastic simulation results, as growing microtubules are much more abundant (see Eq. 10 and Eq. 11). Moreover, the steady-state length distribution in the stochastic simulation (Fig. 1) agrees with the numerical solution (Fig. 3A-3C hollow circles for simulation distribution and red curves for numerical solution of ODEs). The severing activity *k* has profound effects on both growing and shrinking microtubules: increasing severing leads to the tightening of the distribution, reduces the average length and increases the disappearance rate *v*_*s*_*p*_*s*_(0^+^).

**Figure 3.**
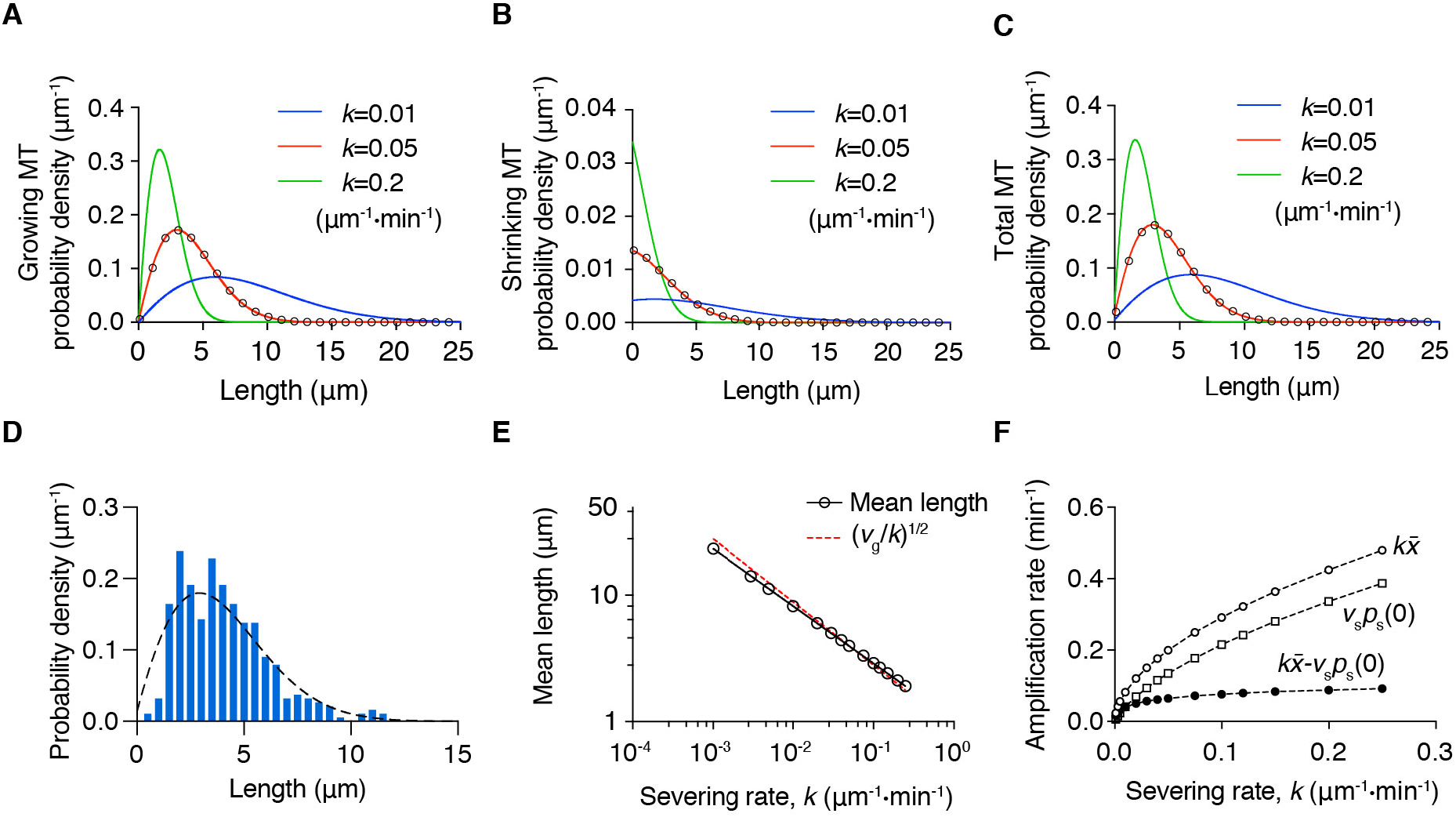
Numerical solution of the steady-state severing model with dynamic instability. Distributions of growing (**A**) and shrinking (**B**) microtubules computed numerically from the ODEs (Eq. 14 and Eq. 15) with boundary conditions (Eq. 22) are plotted for three different severing rates *k*. The dynamic parameters used for the solution are contained in Table 1. More frequent cutting leads to the shortening and compaction of length distribution. **C** The total microtubule length distribution, which is the sum of distributions in **A** and **B**. The steady-state distributions from the stochastic simulations (*k*=0.05 μm^−1^⋅min^−1^) are shown as hollow circles in **A**-**C** and agree with the ODE solution. The proportion of shrinking microtubules is much smaller than that of growing microtubules, so the total distribution is similar to the growing one. **D** Comparison of the experimental length distribution (blue histogram, (19), with the predicted length distribution (dashed line, *k* = 0.05 μm^−1^⋅min^−1^). Both distributions have a mean length of approximately 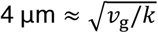. **E** Log-log plot of the mean length as a function of the severing rate. The hollow circles are the mean length obtained from the numerical solution. Black line is the linear regression of log *k* and 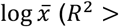 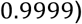. Red dashed line indicates (*v*/*k*)^1/2^ of the no-catastrophe model case where *v* = *v*_*g*_ = 0.79 μm/min. **F** Severing rate *k* versus the amplification rate 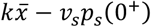 (black circles), the average number of cuts on a single microtubule 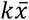 (hollow circles), and the microtubule disappearance rate *v*_*s*_*p*_*s*_(0^+^) (squares). These three functions all increase with severing activity, but the amplification rate quickly reaches a plateau.

The model is in good agreement with the experimental results (19). With a severing rate *k* = 0.05 μm^−1^⋅min^−1^, similar to that measured experimentally, the mean length is approximately 4 μm, and the predicted length distribution (dashed-line in Fig. 3D) resembles the observed one (Figure 3D histogram). We have not attempted to comprehensively test the model against experiments as several of the experimental parameters are difficult to measure precisely. For example, severing can be difficult to distinguish from catastrophe and the length distribution is difficult to measure when microtubule fragments are released from the surface. Furthermore, the theory makes simplifying assumptions such as no minus-end growth and the new plus ends always starting in the shrinking state; the experimental results show that these assumptions only hold approximately. Nevertheless, the good agreement between the measured and predicted steady-state length distributions seen in Figure 3D is a strong qualitative support for the model.

Unexpectedly, when investigating the impact of severing activity on the mean length, we found that the log-log plot is highly linear, with a slope of ≈ −0.45 when the mean length (on the y-axis) is plotted against the severing rate (Fig. 3E black circles and line). This power-law is close to the 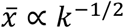 relation in the no-catastrophe model (Eq. 31). Further comparison demonstrates that in the presence of dynamic instability, the mean length is close to the no-catastrophe case ((*v*/*k*)^1/2^, Fig. 3E red dashed line from Eq. 31). This suggests that the addition of dynamic instability in this regime has a relatively small impact on the average length, though we do not have a good explanation for the deviation from a slope of −1/2.

At steady state, the total microtubule mass and number increase exponentially and the amplification rate, 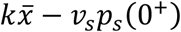, is determined by the competition between the speed of generating new microtubules by cutting and the disappearing of old microtubules (Eq. 4 and 5). With increasing severing rate, the average number of cuts on a single microtubule 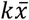 as well as the disappearance rate *v*_*s*_*p*_*s*_(0^+^) increase (Fig. 3F hollow circles and hollow squares). The amplification rate also increases with the severing rate (Fig. 3F solid circles), but with a lower slope at higher severing activity. These results demonstrate that faster severing can lead to faster expansion of the microtubule network, but the effect quickly saturates with increasing cutting rate due to the shortening of lengths and higher probability of losing microtubules.

### Microtubule growth rate is a key regulator of length distribution and amplification rate

Motivated by the close resemblance of the mean length in the full dynamic case and the no-catastrophe model, we next examined the effect of growth rate *v*_*g*_ by solving the length distribution at a severing rate *k* = 0.05 μm^−1^⋅min^−1^ with various growth rates, ranging from that measured with tubulin alone to that measured in high concentrations of the polymerase XMAP215 (35). Similar to the no-catastrophe case, higher growth rate leads to longer lengths of microtubules and broader length distributions (Fig. 4A-B). Interestingly, the proportion of shrinking microtubules also increases with growth rate, while the disappearance probability *p*_*s*_(0^+^) is insensitive to this change (Fig. 4B). The mean length and polymerization rate also showed a power-law relation with a power of ∼0.45 (Fig.4C black circles and line). Similar to the earlier findings, the mean length can also be well approximated by (*v*_*g*_/*k*)^1/2^ (red dashed line in Fig. 4C). Owing to the combination of longer mean length, which increases the average number of cuts per microtubule, and the invariance of disappearance probability, the amplification rate 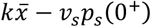 shows a strong increase with the polymerization rate (Fig. 4D solid circle). This large increase of the amplification rate with growth rate compared to the small increase with the severing rate (Fig. 3F) shows that modulation of the amplification rate is more effectively achieved by changing the growth rate rather the severing rate, even though they both strongly affect the mean length. Thus, the polymerization rate substantially affects both the length and amplification rate of the microtubule network.

**Figure 4.**
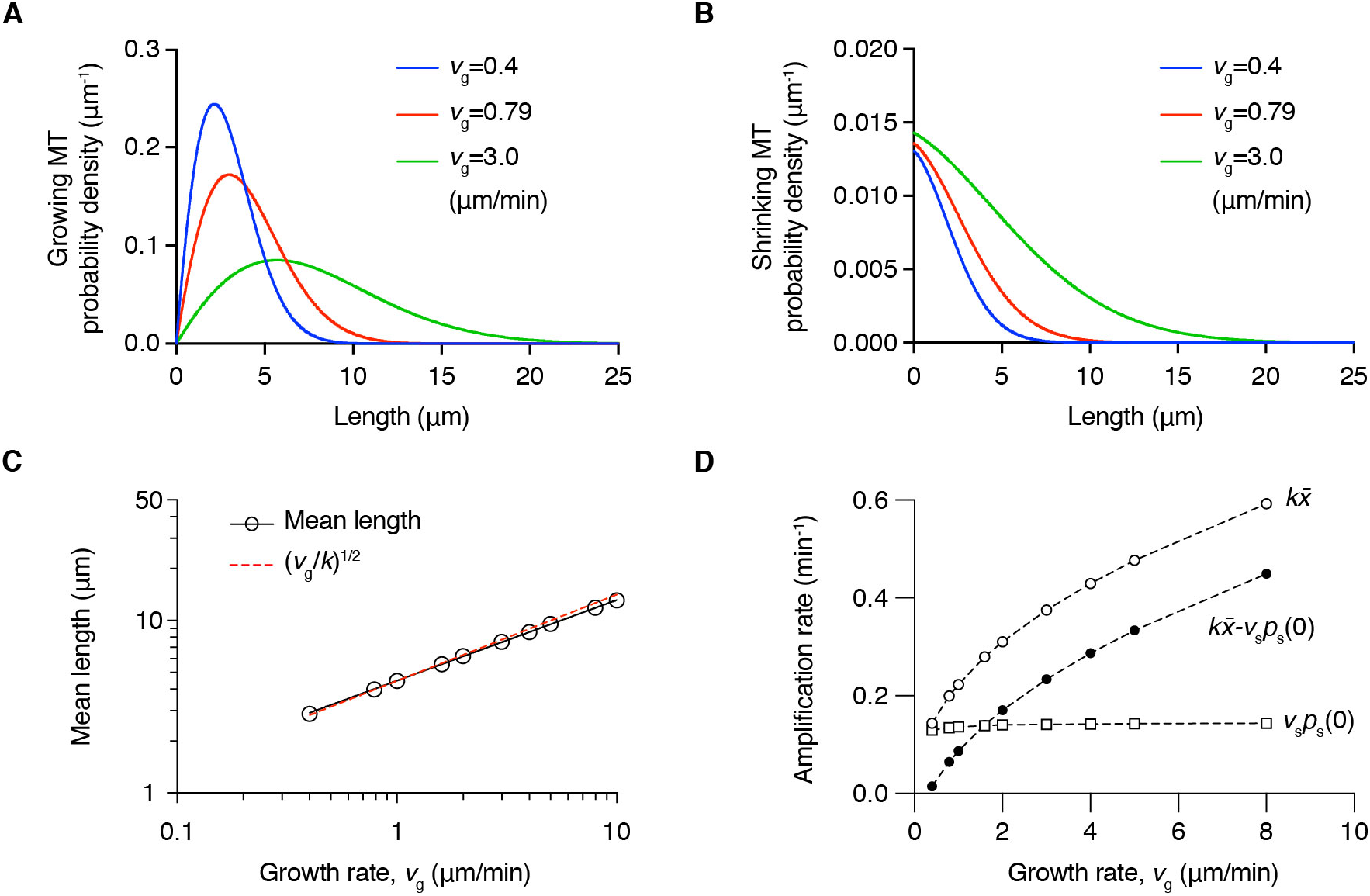
Effect of growth rate on the steady-state length distribution. The growing (**A**) and shrinking (**B**) microtubule length distributions with different growth rates *v*_*g*_. The severing rate *k* is 0.05 μm^−1^⋅min^−1^. Faster polymerization rates increase the mean length and broaden the distribution. The disappearance probability *p*_*s*_(0^+^) varies little with the growth rate (see the *y*-intercept in **B**). **C** Steady-state mean length and growth rate from the numerical solution (circles) shows a power-law relation with a slope close to ½ (red dashed line). **D** Amplification rate 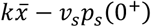 (black circles), average number of cuts on a single microtubule 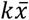 (hollow circles), and microtubule disappearance rate *v*_*s*_*p*_*s*_(0^+^) (squares) versus the growth rate *v*_*g*_. The mean length increases quickly with growth rate while the microtubule disappearance rate is much less affected. The amplification rate, 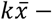 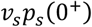, therefore increases with faster growth, mainly due to the longer mean length and the increasing number of cuts on a single microtubule.

### Steady-state length distribution is insensitive to other dynamic parameters

Next, we explored how the other dynamic parameters (shrinkage rate *v*_*s*_, catastrophe *f*_*gs*_ and rescue frequency *f*_*sg*_) affect the steady-state length distribution. Constrained by the unbounded growth criterion, which is essential for increasing microtubule mass with severing, the rescue frequency has a lower bound and the shrinkage rate and catastrophe frequency have upper bounds. We found that these parameters have a comparably small effect on the steady-state mean lengths, even when varied over physiologically relevant ranges attained in the presence of various MAPs (e.g. EB1 increases catastrophe to ∼1 min^−1^(35), CLASP increases rescues up to 10 min^−1^ (36), and spastin and TPX2 decreases shrinkage to ∼5 μm/min(37)) (Fig. 5A-C). In all tested conditions, the change on the mean length was within 0.5 μm (∼10%).

**Figure 5.**
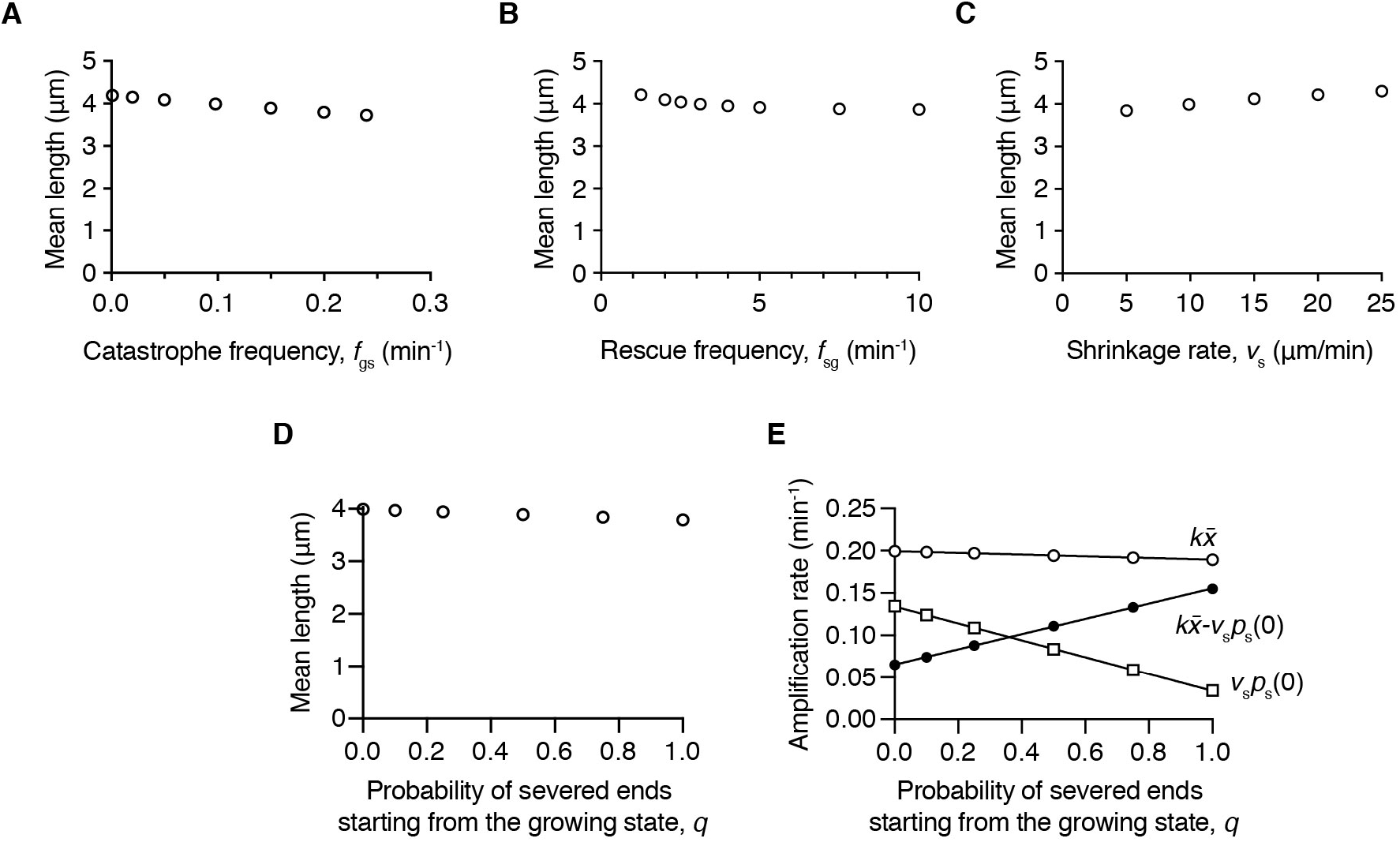
Effects of dynamic parameters on the steady-state mean length. Higher catastrophe frequency (**A)** and rescue frequency (**B)** shorten the mean length, while increasing the shrinkage rate leads to a longer average length (**C)**. The mean length change is small. **D** The mean length depends only weakly on the probability that a newly generated plus end starts in the growing state, denoted by *q*. Earlier in this analysis we assumed that *q* = 0. **E** The amplification rate (black circles) increases strongly as the probability that a newly generated plus end starts in the growing state. This increase is mainly due to the decrease in the rate of disappearance of microtubules (open squares).

As one might expect, a higher catastrophe frequency decreases the microtubule length (Fig. 5A). However, higher rescue frequency and lower shrinkage rate actually shorten the steady-state mean length (Fig. 5B and C), which may appear to be counterintuitive at first sight. Examining the length distribution in these cases revealed that this is due to the increasing survival of shorter microtubules. Lower shrinkage rate leads to an increase in shorter microtubules, both growing and shrinking ones (Fig. S3A and B). As opposed to the growth velocity, varying the shrinkage velocity has a relatively small impact on the growing microtubules (Fig. S3A), but changes the shrinking microtubule distribution more prominently (Fig. S3B). On the other hand, the rescue frequency modulates both the growing and shrinking microtubule distributions: a high rescue frequency increases the proportion of short growing microtubules (Fig.S3C) while it decreases the amount of short shrinking ones (Fig. S3D). Due to the dominance of growing microtubules (Eq. 10 and Eq. 11), increasing the rescue frequency leads to an overall shorter average length (Fig. 5B). Despite the mean length depending only weakly on the shrinkage rate and rescue frequency, they have a more pronounced effect on the amplification rate by modulating the microtubule disappearance rate. In conclusion, these results show that promoting rescue and slowing down shrinkage can decrease the microtubule disappearance rate and lead to a faster amplification with little effect on length.

### Effect of stabilizing newly generated plus ends

All the modeling thus far has assumed that the new plus ends generated by severing start in the shrinking phase. This assumption is based on *in vivo* and *in vitro* experimental results showing that ∼85% of new plus ends are shrinking (see Methods). This fraction can be regulated by plus-end binding proteins such as CLASPs(27) in cells.

To investigate how the state of the newly created plus ends affects the microtubule amount and length distribution, we extended Eq. 1 and 2 to include the probability (denoted by *q*) that a newly generated end starts in the growing phase immediately after severing and obtained the new time evolution equations (Eq. 23 and Eq. 24). The summation of these two equations is independent of *q* and thus the temporal solutions of microtubule number and mass are unchanged, and both increase exponentially with time when the distributions reach a steady-state (Eq. 4 and Eq. 5).

The steady-state length distribution can be solved numerically with the same approach (Eq. 25 to Eq. 27). Increasing the probability *q* is analogous to rescue promotion and leads to a slight shortening of growing microtubules (Fig. S2E, the distributions shift left with higher *q*), and decreases the proportion of shrinking microtubules (Fig. S2F).

The steady-state mean length is only weakly perturbed by the stabilization of the newly generated plus ends (Fig. 5D) (using the dynamic parameters in Table 1). Intriguingly, the disappearance rate of microtubules (*v*_*s*_*p*_*s*_(0^+^) decreases almost linearly with *q* (Fig. 5E, hollow squares); therefore, the stabilization of newly created plus ends (*q*) almost linearly increases the amplification rate (Fig. 5E black circles). Thus, stabilization of severed ends can be a potent method to generate microtubules with a minor change in their length.

### Comparison to a system with a steady-state number of microtubules

Previous work by Tindemans & Mulder also investigated the impact of severing on the steady-state length distribution. In their scenario, a constant spontaneous nucleation rate balances the loss of microtubules from shrinkage, leading to a constant number of microtubules(26). This corresponds to the bounded growth condition in the Dogterom & Leibler model (*f*_*sg*_*v*_*g*_ − *f*_*gs*_*v*_*s*_ < 0)(21). In contrast, we consider the scenario where the number of microtubules is increasing, corresponding to the unbounded growth condition (*f*_*sg*_*v*_*g*_ − *f*_*gs*_*v*_*s*_ > 0). These two scenarios give rise to important differences between the length distributions. First, in the Tindemans & Mulder scenario, the steady-state microtubule number is independent of the severing rate, while we show that the number increases exponentially with a rate that increases with the severing rate (Fig. 3F). Second, in the Tindemans & Mulder scenario, spontaneous nucleation and the bounded growth condition together cause a high percentage of very short microtubules, while in our scenario the proportion of short microtubules is small due to a higher survivability of longer microtubules (Fig. 3C). Third, in the Tindemans & Mulder scenario, the ratio of the total number of growing and shrinking microtubules is equal to the ratio of the shrinkage and growth rates, while we found that severing introduces an additional bias towards growing microtubules (Eq. 10 and 11), and thus gives rise to an overall increase in the total microtubule mass and number. Last, the power-law relations between the mean length and the severing or growth rate arise only in the scenario considered here (Fig. 3E and 4C). Thus, the constraint of constant microtubule number and the presence of spontaneous nucleation profoundly affect the length distribution and total microtubule number.

In the Tindemans & Mulder scenario, spontaneous nucleation is essential to compensate for the loss of shrinking microtubules. This scenario, which may be important in plant cells, may not be as relevant in animal cells, where spontaneous nucleation of microtubules (away from the centrosome) is normally rare (38, 39). Our results suggest that severing can also serve as a nucleation-like mechanism that rapidly increases the production of microtubules in the unbounded growth regime, regardless of the spontaneous nucleation rate. Indeed, if there is an exponential increase in new microtubules by severing and regrowth, existing nuclei or spontaneous nucleation will make decreasing contributions to the total number of new microtubules. However, if the amplification is autocatalytic, other cellular mechanisms will be required to stop this activity before free tubulin is depleted.

### The similarity between models with and without dynamic instability in the presence of severing

Microtubule dynamics measurements in various systems such as sea urchin and *Xenopus* egg extracts(40, 41), budding yeast (42), *C. elegans* (43), and tissue culture cells(44–46) have shown that the *in vivo* dynamic parameters are highly diverse across different species, cell lines and cell cycle stages. Growth and shrinkage rates can span from ∼0.3-20 μm/min and ∼5-50 μm/min respectively. Wide ranges also exist for cellular catastrophe (∼0.3-10 min^−1^) and rescue frequency (< 0.1-20 min^−1^), and subsets of highly stable microtubules with little turnover have been observed in tissue culture cells and neurons(47, 48). While our simulations have not covered this entire range, they have shown some general principles of how severing influences microtubule length and number. A surprising finding is that the steady-state mean length can be well approximated by the no-catastrophe simplified model (Fig. 3E and 4C), implying that in the presence of severing, the effect of dynamic instability on the length distribution is fairly small. This phenomenon can be understood by the fact that the total growing time of microtubules is much longer than the shrinking time, and thus approximates the simplified model where no shrinking microtubules are present. On average, the growing time before catastrophe is 1/*f*_*gs*_ and the shrinking time before rescue is 1/*f*_*sg*_. The microtubules with a length shorter than *v*_*s*_/*f*_*sg*_ will shrink and disappear faster, leading to an even shorter mean shrinking lifetime.

A final point is that microtubule dynamics with severing is a non-ergodic system: the ratio of average growing and shrinking lifetimes is higher than the proportion of growing and shrinking microtubule numbers. This is because the short shrinking microtubules vanish rapidly. In the unbounded growth condition, where the rescue frequency is high and catastrophe frequency is low, a single microtubule is thus predominantly in the growing phase, and this can explain why the overall effect of severing on length resembles the no-catastrophe model even when dynamic instability is present. Thus, the impact of dynamic instability on the steady-state length distribution is fairly small and the dynamics can be well approximated by a model that considers only microtubule polymerization and severing.

## Conclusions

We have explored how microtubule dynamics affects the length distribution and the amplification rate of microtubule number and mass in the presence of severing by mathematical modeling. Unexpectedly, dynamic instability has a small impact on the steady-state length, at least over observed microtubule dynamics parameters. The microtubule length is mainly governed by the polymerization and severing rates, and can be well-approximated by a no-catastrophe simplified model, which we have solved analytically. Rescue frequency, catastrophe frequency, shrinkage rate and the probability that newly severed ends start in the growing phase perturb the length distribution only weakly, but have a more profound impact on the amplification rate. Comparison with previous experimental measurements provides strong qualitative support for our mathematical model, despite the simplifications such as omitting the minus end dynamics and assuming severing as a single-step instantaneous process.

Cellular microtubule lengths are controlled by various machineries(49–51) including depolymerases(52–54), polymerases(55, 56), the centrosome(57), severases(58) and motors(59, 60). Our theoretical analysis shows that microtubule severing, in addition to shortening microtubules, also makes the microtubule length distribution more uniform. For example, the coefficient of variation of microtubule lengths (SD/mean) is ∼0.58 for the parameters used in Table 1 (see Figure 3C), compared to 1, which is for the exponential distribution that solves the Dogterom & Leibler model under the bounded growth condition in the absence of severing. In this respect, severing has a similar functional consequence to the length-dependent depolymerase kinesin-8(52), which also tightens the length distribution of dynamic microtubules(53, 61). Our results also demonstrate that spastin has an effective nucleation activity: the exponential increase in microtubules is a consequence of the microtubule-dependence of severing, and in this respect, nucleation by severases is analogous to the explosive nucleation by augmin, which nucleates new microtubules from the sides of extant microtubules (51, 62). Thus, our analysis provides a quantitative understanding of how severing and dynamics can jointly regulate the morphology of microtubule networks.

## Author contributions

Y.W.-K. and J.H. formulated the model. Y.W.-K. performed the experimental measurements and numerically integrated the stochastic differential equations at steady-state. O.T. solved the differential equations of the simplified model and performed the stochastic simulations. All authors wrote the manuscript.

## Acknowledgement

We thank the current and previous Howard lab members for the feedbacks on this work. Y.-W.K. is supported by a fellowship from the Ministry of Education in Taiwan. This work was supported by NIH Grants R01 GM110386 (to J.H.) and DP1 M110065 (to J.H.).

## Appendix

## Analytical solution of simplified no-catastrophe model

The master equation of the simplified no-catastrophe model reduces to the following integro-differential equation at steady state:

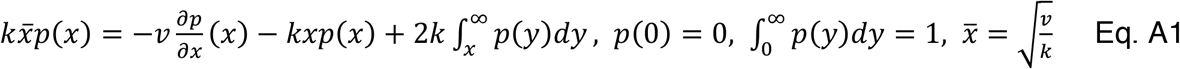

First, we differentiate with respect to *x* to obtain an equivalent second-order differential equation:

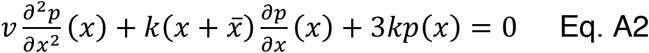

Using the normalization condition on *p*(*x*), (Eq. A1) we can derive a boundary condition for 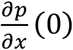 that is consistent with the differential equation:

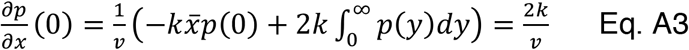

Using the substitution 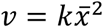, the differential equation problem becomes:

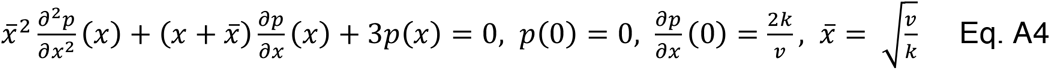

Next, we perform a change of variable to center the distribution around the mean. Let 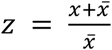, the problem becomes:

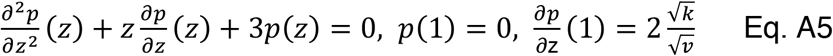

By performing the substitution 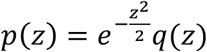, the differential equation becomes:

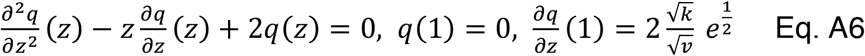

The second order ODE is recognized as the Hermite differential equation for the specific case where λ = 2. The convergent solutions of the Hermite equation are known as the Hermite polynomials, which were first described by Pierre-Simon de Laplace and later by Pafnuty Chebyshev and Charles Hermite in the 1800s. Since we are seeking well-behaved solutions, we require that *q*(*Z*) be polynomially bounded and the solution for *q*(*Z*) corresponds to the second-order Hermite polynomial:

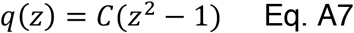

where *C* is a constant that can be determined by the boundary condition. One notable application of Hermite polynomials in physics is the quantum harmonic oscillator, where they give rise to the eigenstates of the Schrödinger equation(34). The solution for Eq. A6 is also found when solving the wave function of the second excited state in quantum harmonic oscillator system (vibrational quantum number *v* = 2 for a one-dimensional molecular vibrational system).

To satisfy the boundary conditions, we set 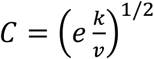. Reverting the substitutions, the final solution becomes:

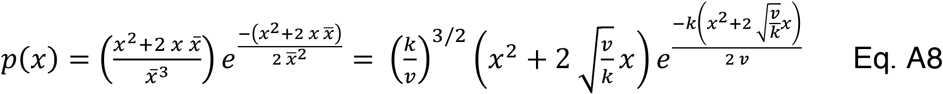

---

**Supplementary Figure 1.**
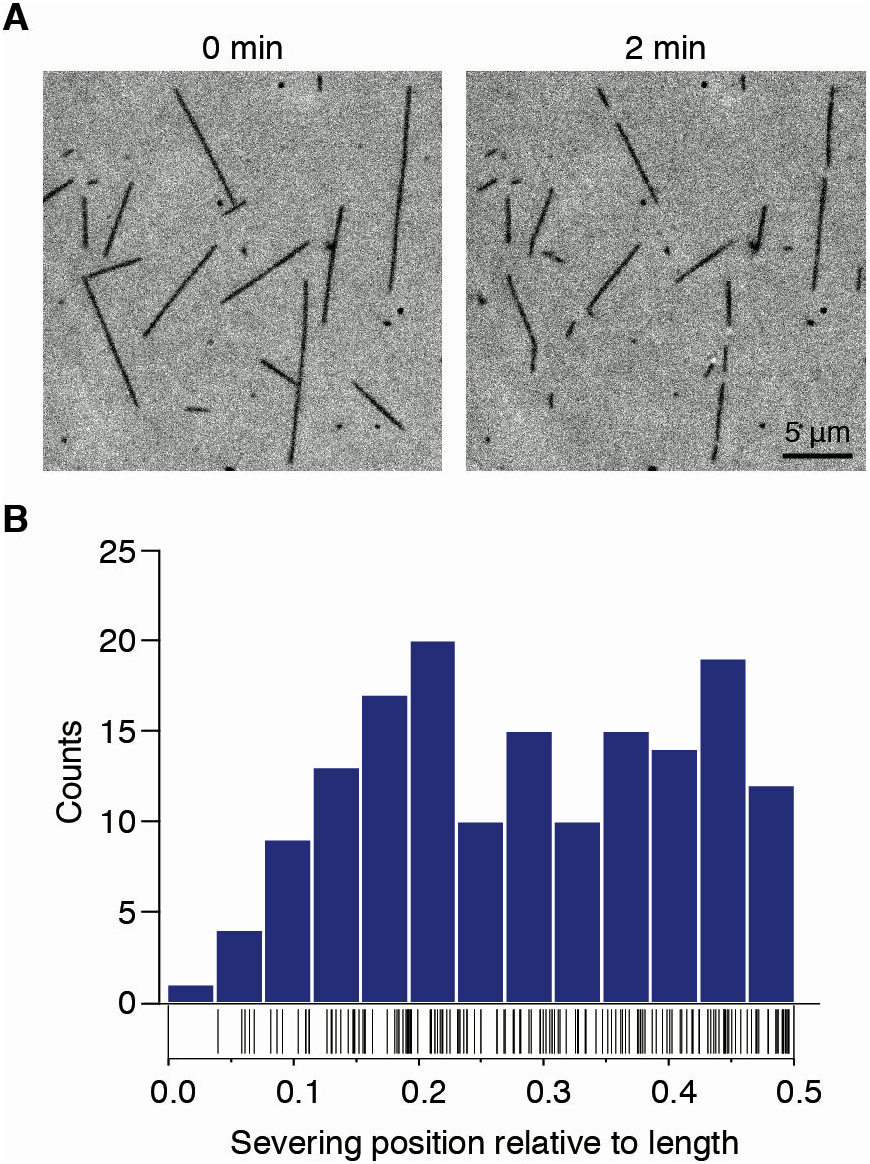
Severing positions on stabilized microtubules. **A** Example of GMPCPP-stabilized microtubules severed by *Drosophila* spastin. Breakages of microtubules are visible with interference reflection microscopy (IRM). **B** Distribution of severing position along microtubule length showed as histogram and rug plot. The severing position is quantified by measuring the shorter fragment length divided by the full length before a cut occurred. The lower frequency near the tip (<0.1) results from the difficulty of detecting short fragments limited by the optical resolution. The uniformity of the severing position is tested using a chi-squared test that excludes the first three bins. The test results (χ^2^=7.34, p-value = 0.60, degrees of freedom=9) suggest that the experimental distribution is consistent with a uniform distribution. The total number of measurements (N=159) were collected from duplicate experiments.

**Supplementary Figure 2.**
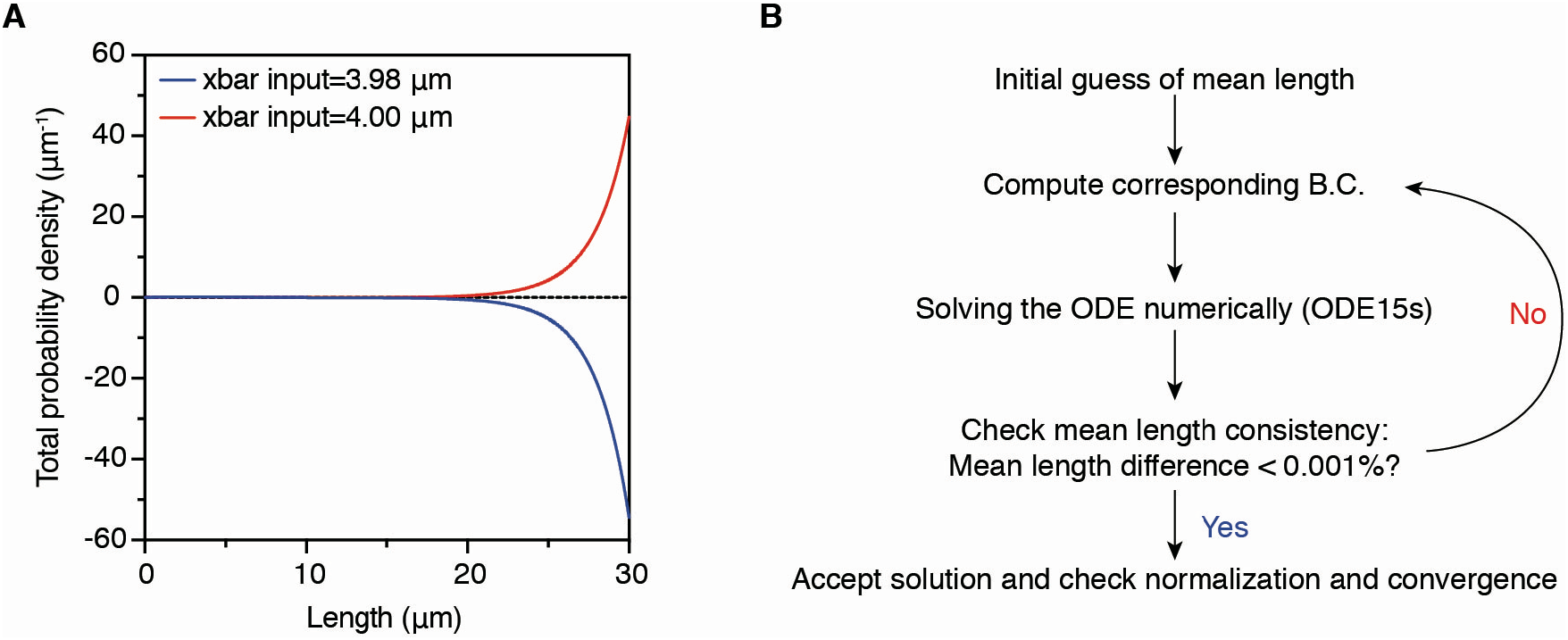
Numerical solution of microtubule length distribution. **A** The numerical integration results diverge with opposite sign when input 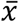 deviates from the true mean length, and the direction depends on whether it is an over- or under-estimation. The dynamic parameters used were described in Table 1, with a severing rate of 0.05 μm^−1^min^−1^. The true mean length is 3.991 μm in this condition (Fig. 3A-3C, red curves for the converge and self-consistent solution). **B** Iterative procedure for solving the steady-state length distribution numerically. Normalization error is smaller than 0.001.

**Supplementary Figure 3.**
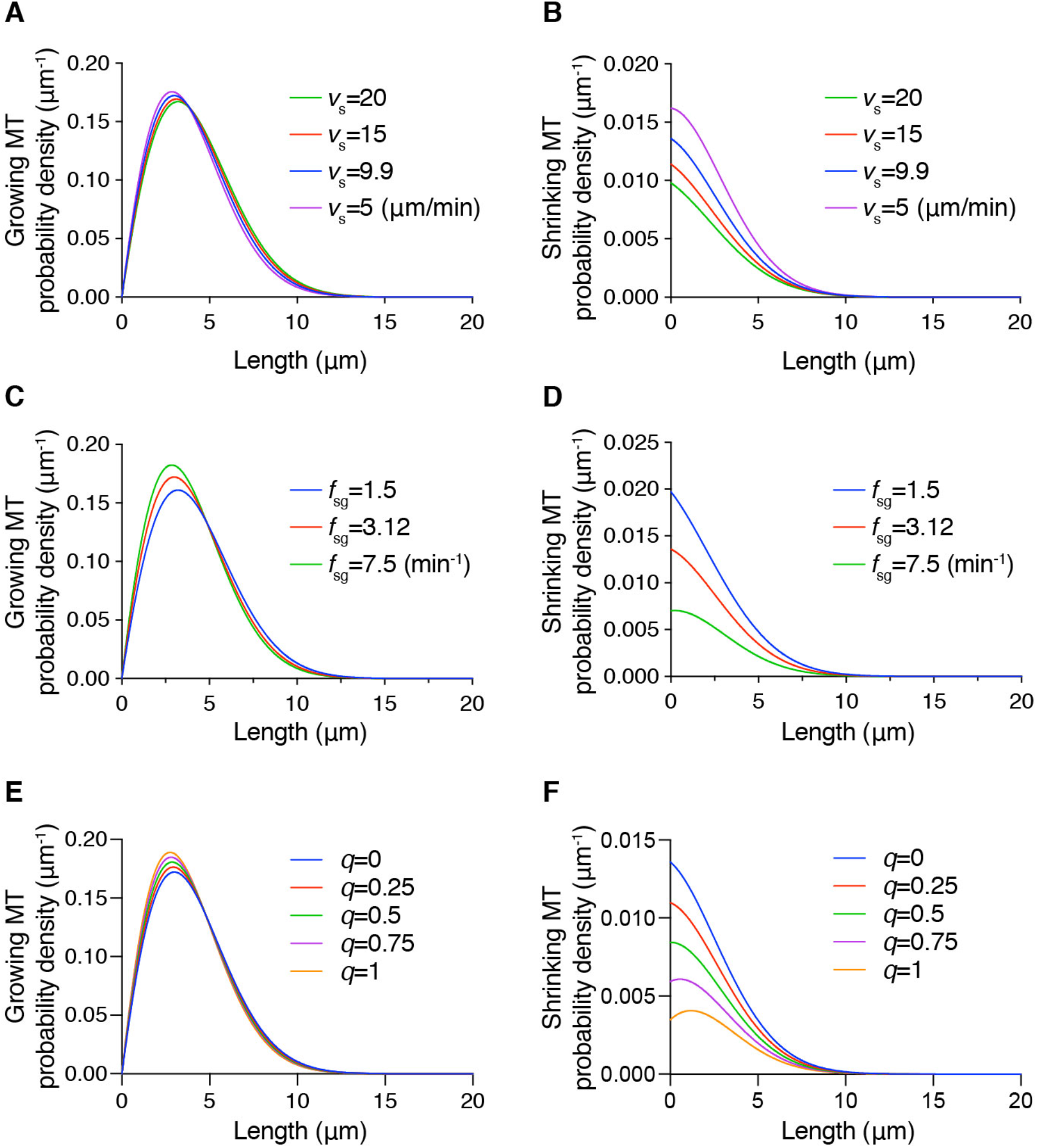
Steady-state length distribution with respect to different dynamic parameters. **A,B** Effect of shrinkage rate *v*_*s*_ on growing and shrinking microtubule distribution. The growing probability distribution is almost unperturbed while the shorter shrinking microtubules probability is more affected. **C,D** Growing and shrinking microtubule distribution for different rescue frequencies. Promotion of rescue has an opposite effect on growing and shrinking distribution: it increases the amount of short growing microtubules but decreases the amount of short shrinking ones. **E,F** Steady-state length distribution with different probability of new plus ends immediately starting in the growing state after cut (denoted by *q*). The effect of *q* is similar to rescue and has a strong effect on the microtubule disappearance probability *p*_*s*_(0^+^). The solutions are solved with a severing rate of 0.05 μm^−1^min^−1^.

